# Proteome structuring of crown-of-thorns starfish

**DOI:** 10.1101/2024.08.25.609624

**Authors:** Zhu Yunchi, Lu Zuhong

**Affiliations:** State Key Laboratory of Digital Medical Engineering, School of Biological Science and Medical Engineering, Southeast University, Nanjing, China

**Keywords:** proteome structuring, crown-of-thorns starfish, Acanthaster planci, ColabFold, structural bioinformatics

## Abstract

This data report presents the proteome structuring of the crown-of-thorns starfish (COTS, *Acanthaster planci*) using the ColabFold system. The resulting dataset includes 31,743 predicted protein structures, covering 60.4% of residues with confident predictions and 35.5% with very high confidence. A preliminary structural bioinformatics analysis was conducted using post-AlphaFold methods, including fast structure clustering, ligand transplanting, and structure-based Gene Ontology (GO) annotation, which identified potential functions for previously uncharacterized proteins. These findings contribute to a deeper understanding of COTS biology and provide valuable data for developing biocontrol methods to mitigate the impact of COTS outbreaks on coral reefs.

## 1 Introduction

The crown-of-thorns starfish (COTS, *Acanthaster planci*) is a highly fecund predator of reef-building corals throughout the Indo-Pacific region (1). COTS population outbreaks cause significant damage to coral reefs, the living environment for more than 30% of marine animals and plants (2), leading to a loss of coral cover and biodiversity. Scientists have sequenced the COTS genome (1), which provides a wealth of information on the genetic basis of COTS biology. By identifying specific genes and proteins involved in these behaviors, scientists are able to gain a deeper understanding of their reproductive strategies and the factors contributing to outbreaks, so as to develop targeted biocontrol methods such as peptide mimetics to disrupt COTS aggregation. However, the function annotation of COTS proteome turns out to be incomplete, with over 20% of proteins being annotated as “uncharacterized”. Traditional sequence-based annotation methods may be insufficient for fully resolving genomes, particularly for non-model organisms. It is commonly recognized that “sequence determines structure, and structure determines function.” If proteome structuring, by which sequences can be transformed into accurate structures in a high-throughput way, is as feasible as genome and transcriptome sequencing, it is believed that such approach could not only substantially aid researchers in complementing and correcting protein annotations, but also pave a new dimension for protein data mining (3).

This vision is going to be realized with the help of the booming artificial intelligence (AI) technology. AI-based protein structure prediction systems represented by RoseTTAFold (4) and AlphaFold2 (5) have brought the dawn of the high-throughput era of structural proteomics. With accuracy not inferior to traditional methods such as x-ray crystallography and cryo-electron microscopy (4), they have overwhelming advantages in cost, efficiency and ease of operation. As of July 2022, the AlphaFold Protein Structure Database (AFDB) boasts open access to a staggering collection of over 200 million protein structures (6), marking a 1000-fold increase compared to the 50-year accumulation of the PDB. Furthermore, owing to collaborative efforts within the open-source community, many optimized versions of AlphaFold2, which are collectively referred to as AlphaFold-like systems have been developed, with ColabFold (7) standing out as a notable example. It significantly reduces the resource demands for protein folding, empowering more researchers to engage in personalized structure predictions and expanding the overall data scale. Up to now, AlphaFold-like systems present to be the most robust tools for high-throughput proteome structuring (8), facilitating a diverse array of research endeavors. Notable examples include ColabFold proteome CP-8382 from Southeast University (2), structural proteome of *Sphagnum divinum* from Oak Ridge National Laboratory (9), AlphaFold proteome of *Mnemiopsis leidyi* from National Institutes of Health (10), etc. These explorations are progressively expanding the protein structure universe and enabling new insights into protein function and biology.

Here we present the proteome structuring of COTS. Deploying ColabFold in the Big Data Computing Center at Southeast University, we predicted 31,743 protein structures. The resulting dataset covers 60.4% of residues with a confident prediction and 35.5% with very high confidence. We also performed a preliminary structural bioinformatics analysis using several post-AlphaFold methods, including fast structure clustering, ligand transplanting and structure-based Gene Ontology (GO) annotation.

## 2 Materials and methods

### 2.1 Proteome structuring

The NCBI RefSeq of *Acanthaster planci* (GCF_001949145.1) was used as sequence source. Protein sequences were downloaded and filtered, discarding those exceeding 2,550 aa to accommodate the upper limit of GPU memory. Multiple sequence alignments (MSA) generation was conducted locally (*colabfold_search*), then MSAs in A3M format were uploaded to ColabFold 1.5.2 on the NVIDIA Tesla V100 cluster at the Big Data Computing Center of Southeast University. The parameters of ColabFold were set to *–amber, –num-recycle 3, –use-gpu-relax, --zip, --num-relax 1*.

During the structure prediction process, the MineProt (11) toolkit (*colabfold/import*.*sh --name-mode 0 --zip --relax*) was periodically executed to process predicted proteins. This included selection of best structure models with highest predicted local distance difference test (pLDDT) scores, generation of CIF files, and storage of model scores in JSON format.

### 2.2 Structure alignment and clustering

Foldseek (12) was employed for high-throughput structure alignment clustering. Predicted structures were aligned to the AlphaFold Clusters (13) using *easy-search –e 0*.*01 -s 7*.*5*, and were clustered using *easy-cluster -c 0*.*9 -e 0*.*01 --min-seq-id 0*.*5*.

### 2.3 Structure-based function annotation

The representative protein structures identified by Foldseek clustering got function annotation. AlphaFill (14) 2.0.0 with PDB-REDO databank (15) was used to enrich them with ligands and cofactors, and DeepFRI (16) was used for GO annotation.

Uncharacterized proteins with average cluster pLDDT > 70 were selected for GO enrichment analysis by the aid of clusterProfiler (17), where GO annotations with DeepFRI score lower than 0.5 were pre-filtered. The parameters were *pvalueCutoff = 0*.*05, pAdjustMethod = ‘fdr’, qvalueCutoff = 0*.*2*.

## 3 Results

The resulting dataset contains 31,743 protein structures, of which 6,338 were tagged with “uncharacterized” previously. The model confidence distribution is demonstrated in **Fig 1.A**. 60.4% of residues have a pLDDT larger than 70, and 35.5% have a very high pLDDT over 90. A total of 20,419 protein structures attain an average pLDDT greater than 70, the commonly recognized benchmark of confident model (3).

**Fig 1.**
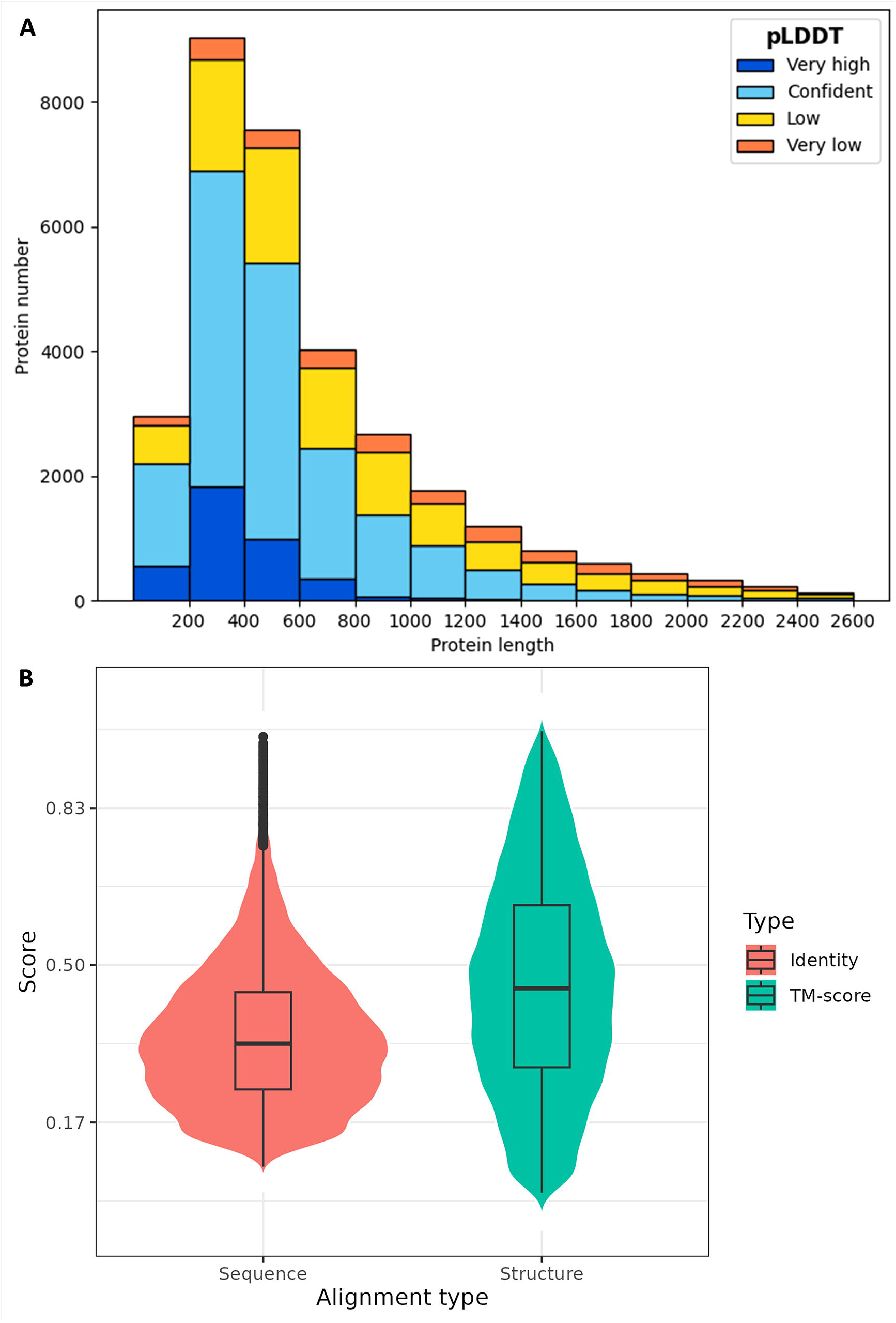
Statistics of COTS proteome structuring. **(A)** Distribution of model confidence against protein length. Horizontal axis is protein length and vertical axis is protein number. Model confidence calculated by pLDDT is color-coded. Very high: pLDDT > 90; confident: 90 > pLDDT > 70; low: 70 > pLDDT > 50; very low: pLDDT < 50. **(B)** Violin plot of structure alignment between COTS structural proteome and AlphaFold Clusters. Horizontal axis is alignment type, for Foldseek performs structure alignment while also calculating sequence alignment. Vertical axis is alignment score, with identity corresponding to sequence alignment and TM-score to structure alignment.

Structure alignment was made between COTS structural proteome and AlphaFold Clusters, the Foldseek clustered AFDB. It should be mentioned that Foldseek hold a comparable position in structural bioinformatics to that of BLAST in sequential bioinformatics, for they both enable the feasibility of large-scale searches in their fields. With its help, scientists have successfully clustered all of the structures in the AFDB into 2.3 million clusters, rendering localized operations of AFDB practicable. As illustrated in **Fig 1.B**, the overall protein structural similarity between COTS and known species within the AFDB is found to be moderate to low, with more than half of alignments exhibiting TM-scores (*qtmscore* in **Table S1**) below 0.5 (18). The majority of sequence identities (*fident* in **Table S1**) between these alignments are even lower, aligning with the notion that protein structure is more conserved than sequence. The phenomenon of low similarity may be attributed to AFDB’s limited inclusion of *Acanthaster planci* as well as its closely related species, for the number of *Acanthaster* protein structures does not surpass 100 in the database. Therefore, the predicted COTS structural proteome from this work can currently serve as a starfish-specific extension of AFDB in view of its generally confident model scores.

Foldseek clustering of the COTS dataset was performed with reference to the construction process of AlphaFold Clusters, and 16,896 structural clusters were generated (**Table S2**). 192 uncharacterized proteins were found to have high structural similarity with annotated proteins, as listed in **Table S3**. They are probably remote homologs hidden by sequential difference. Furthermore, there are totally 3,836 clusters that consist solely of uncharacterized proteins, which can be designated as uncharacterized clusters (**Table S4**). Only 1,045 of them are non-singleton clusters (13), and 1,138 of them have average pLDDT > 70.

Representative protein of each cluster got structure-based function annotation, including ligand transplanting by AlphaFill (**Table S5**) and GO annotation by DeepFRI (**Table S6**). Several annotation results are visualized in **Fig 2**.

**Fig 2.**
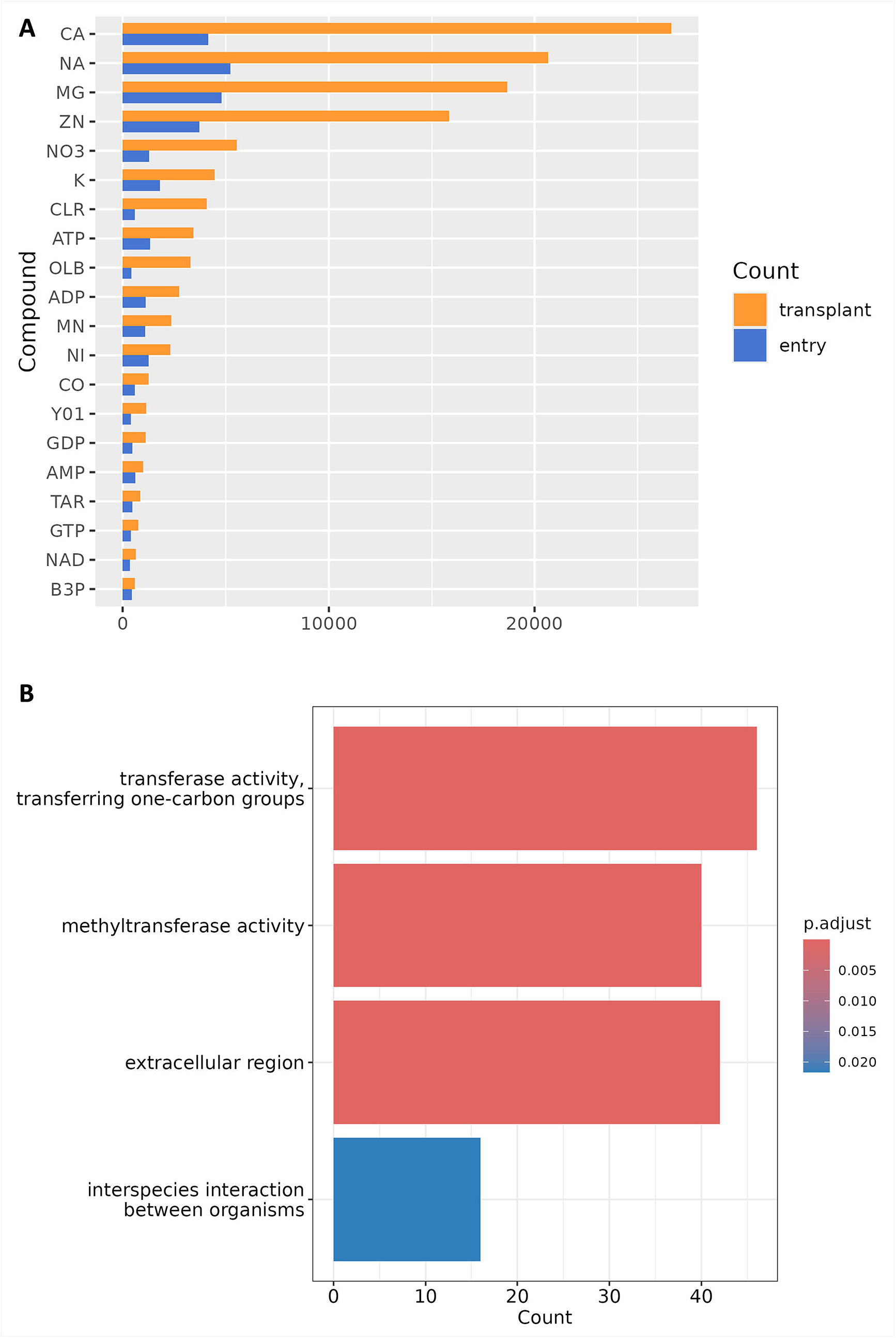
Structure-based function annotation of COTS structural proteome. **(A)** Statistics of top-20 compounds with largest number of transplants. Horizontal axis is number of entries and transplants, and vertical axis is PDBe ligand code. CA: Calcium(^2+^) ion; NA: Sodium(^1+^) ion; MG: Magnesium(^2+^) ion; ZN: Zinc(^2+^) ion; NO3: Nitrate ion; K: Potassium(^1+^) ion; CLR: Cholesterol; ATP: Adenosine triphosphate; OLB: 1-Oleoyl-sn-glycerol; ADP: Adenosine diphosphate; MN: Manganese(^2+^) ion; NI: Nickel(^2+^) ion; CO: Cobalt(^2+^) ion; Y01: Cholesterol hemisuccinate; GDP: Guanosine diphosphate; AMP: Adenosine monophosphate; TAR: d-Tartaric acid; GTP: Guanosine triphosphate; NAD: Nicotinamide adenine dinucleotide; B3P: Bis-Tris propane. **(B)** Bar plot of GO enrichment analysis of representative uncharacterized proteins.

The AlphaFill algorithm identifies experimentally determined protein structures similar to input structure models through sequence alignment, followed by structural comparison to ascertain the positions of ligands and cofactors. Subsequently, these entities are transplanted into structure models, thereby enriching their information content. 10,171 proteins got the “filled” models, and **Fig 2.A** shows the statistics of top-20 compounds with largest number of transplants. Except for cholesterol hemisuccinate (Y01), 19 of them are also present within the top-50 ranked compounds of the entire AlphaFill databank (alphafill.eu). Given that numerous experimentally determined protein structures derive from drug experiments, AlphaFill models may confer additional benefits in supporting the development of marine drugs against COTS outbreaks, particularly in the discovery of small molecule targets.

In the pre-AlphaFold era, DeepFRI is one of the few neural network models for GO annotation that accepts protein structure input. This feature has distinguished it in this era of high-throughput structural proteomics. 13,311 proteins got GO annotation with DeepFRI scores above 0.5, the benchmark of significant prediction (16). Representative proteins of uncharacterized clusters with average pLDDT > 70 were selected for GO enrichment analysis, results of which are demonstrated in **Fig 2.B** and **Table S7**. Three groups of proteins were significantly enriched: one group might be involved in interactions between organisms (GO:0044419), another group is localized to the extracellular region (GO:0005576), and a third group probably have methyltransferase-like activity (GO:0016741 and GO:0008168). These three groups appear to have little overlap, with only XP_022108099.1, XP_022096454.1, and XP_022097466.1 (**Fig S1**) being present in both GO:0044419 and GO:0005576 groups. It should be noted that these three proteins possess signal peptides detectable using SignalIP (19) but no obvious transmembrane domains, which is the feature of secreted proteins (20). Pheromone-like signals are acknowledged as pivotal in the biology of COTS, enabling the regulation of reproductive aggregations, synchronized spawning events (21)(22), foraging behaviors, and escape responses from predators (1). Hence, it might be important for subsequent experimental research as well as meta-analysis to ascertain the presence of these proteins within the COTS exoproteome and to elucidate their potential involvement in conspecific or interspecies communication.

## 4 Discussion

We succeeded to generate a predicted structural proteome of COTS with acceptable confidence. The application of post-AlphaFold structure-centric methodology not only provides evidence for our dataset’s capability of complementing AFDB, but also enhances existing COTS protein annotation. It is expected for the COTS structural proteome to deepen our understanding of COTS biology and facilitating biocontrol method development, not to mention that these protein structures can be directly used as raw inputs in various computational biology tasks, obviating the need for researchers to engage in time-consuming AlphaFold2 deployment and de novo structure modelling.

There is no denying that this work has several limitations. First, the report lacks a thorough interpretation of the ligand transplanting results. The software ecosystem of cheminformatics is less developed than that of bioinformatics, with even the most fundamental ID conversion tools lacking support for PDB ligand codes utilized by AlphaFill. Chemical information mining from these models will continue to pose a challenge unless there is an improvement in the productivity of cheminformatics programmers. Second, the proteome structuring failed to cover proteins too large for our GPU devices to process, a prevalent issue encountered in almost all proteome structuring efforts. It is proposed that reducing computational costs be recognized as another significant direction for the development of AlphaFold-like systems, following the improvement of accuracy and throughput. Last but not least, our COTS structural proteome is entirely a “dry lab” product, necessitating further utilization and assessment by experienced marine biologists. In summary, joint efforts should be made to apply our dataset to the control of COTS and the protection of coral reefs.

## Data availability statement

The datasets generated for this study can be found in Figshare https://doi.org/10.6084/m9.figshare.26706793.v1 (COTS structural proteome) and https://doi.org/10.6084/m9.figshare.26779318.v1 (AlphaFill models).

## Author Contributions

Y.Z.: experiment, writing, and editing. Z.L.: reviewing and supervision.

## Acknowledgments

We thank the Big Data Computing Center of Southeast University for providing the facility support on the numerical calculations in this article.

