## Supplementary figures and images for "Proteome structuring of crown-of-thorns starfish"

### Fig S1

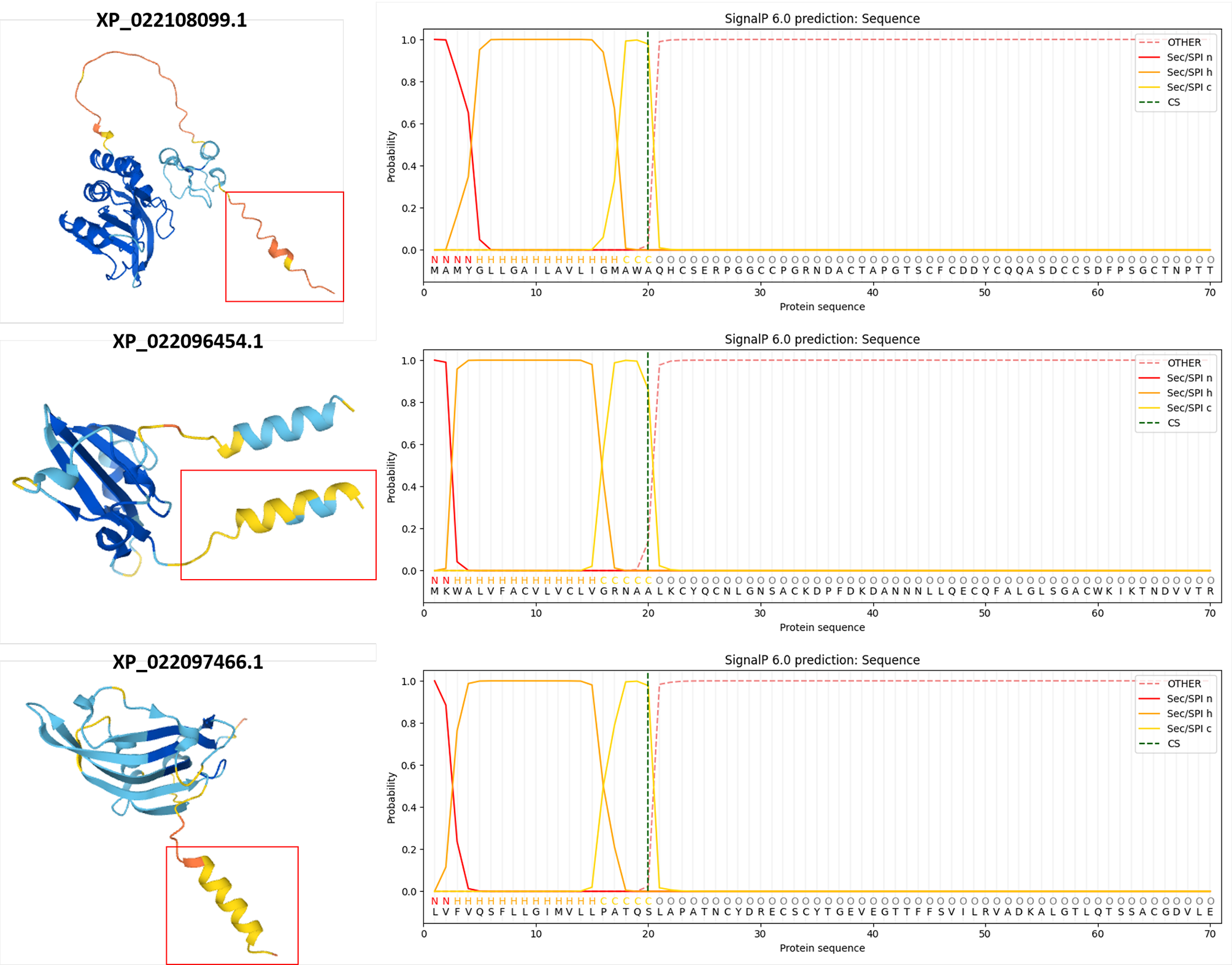
